# Clues on an intraspecific communication system in early plant establishment: The seed-seedling transition from the point of view of a crosstalk between information theory and gene expression

**DOI:** 10.1101/2020.03.12.989269

**Authors:** João Paulo Ribeiro-Oliveira, Lilian E. D. Silveira, Lilian V. A. Pinto, Edvaldo A. A. Silva, Henk W. M. Hilhorst

## Abstract

How much interactivity is there in a seed-seedling transition system? The answer for this question can reveal a key aspect for early plant establishment. Thus, we hypothesize that information entropy is correlated with early plant development because it is directly related to interactions between seed-seed, seed-seedling, and seedling-seedling. To test this hypothesis, we perform an overlapping of classical physiological measurements (embryo protrusion), gene expression in germination *sensu stricto*, water dynamics in germinating seeds and information theory. For a biological model, we used *Solanum lycocarpum* A. St.◻Hil. seeds. This is a Neotropical species with high intra-specific variability in the seed sample. Our finds demonstrate that the dynamic and transient seed-seedling transition system is influenced by the number of individuals (seed or seedling) in the sample, especially at a same physiological stage. In addition, we also discuss that: (*i*) information entropy enables the quantification of system disturbance relative to individuals in the same physiological stage (seed-seed or seedling-seedling), which may be determinant for embryo growth during germination. (*ii*) there is possible intraspecific communication in seed-seedling transition systems formed by germinating seeds with the potential to alter the pattern of embryonic development of the sample. In view of this, we suggest the use of information entropy as a tool for studies of biological systems to clarify the phenomenon of mutual stimulation in the germination process.

## Introduction

The seed-seedling transition is highly defined by parental effects (both maternal and paternal; see Roach & Wulff 1987; Baskin & Baskin 2019). This is because paternal components define the heterogeneity degree among seeds produced along a same reproductive year. The heterogeneity is the basis of intra-specific variability and can be expressed by morpho-physio-genetic components, such as tegument thickness, specific seed mass and/or seed size, as well as embryo vitality and/or seed resilience (e.g., Walter *et al.*, 2016; Imaizumi *et al.*, 2017; Penfield and MacGregor, 2017; Awan *et al.*, 2018; Geshnizjani *et al.*, 2019). These aspects guide the ability of the embryo to capture and process environmental clues in order to manage the intra-seminal growth (see Penfield, 2017; Munz *et al.*, 2017). Thus, we can say that the intra-specific variability in a seed sample is a central component on multiple interactions in a seed-seedling transition system (Labouriau and Valadares, 1976). This interaction is theoretically more intense in artificial environments (e.g., Petri dish, germination plastic boxes and development chamber), where it is expected that populational density will be greater than in natural environments (Labouriau and Valadares, 1976; Ribeiro-Oliveira and Ranal, 2016). Taking this into account, we ask: how can intra-specific variability affect functional understanding of the germination process? A crosstalk between information entropy and fold transcriptional change gives us a safe way to answer that question.

Information entropy was determined by communication science and numerically determines interaction processes of signal transmission and reception in a binary system (Shannon, 1948). The idea of entropy itself was transposed into ecology by means of the index that takes the name of the researcher (Shannon’s entropy or Shannon’s index). In this area, entropy is used to measure population or sample variability (Jost, 2006). In thermobiology science, there are scientists that consider thermodynamic entropy and information entropy as synonyms, which is called the statistical entropy of the thermodynamics of the Boltzmannn theory (Wehrl, 1991). By the way, the little that is currently present in the entropy of germination *sensu stricto* is based on Boltzmann theory (see Dragicevic and Sredojevic, 2011), probably due to the difficulty of determining the information entropy during germination *per se*. However, it has recently been demonstrated that when using water dynamics in germinating seeds at a standard temperature (a highly related character to the germinative metabolic processes), it is possible to calculate the information entropy in a much easier, robust and practical way (Ribeiro-Oliveira, 2015). This entropy measurement reflects discrepancies of delayed and precocious embryo protrusion in a seed-seedling transition system from water dynamics in a germinating seed in the sample. Therefore, this measurement can be useful to clarify the relationship between embryo development and the disorder provoked in the seed-seedling transition system from an ecophysiological point of view. Thus, we hypothesize that information entropy peaks are associated with moments of intense embryo development because they demonstrate an intra-specific communication of individuals in similar physiological stages (seed-seed, seed-seedling or seedling-seedling), demonstrating in practice a mutual stimulation phenomenon. To validate it, we will correlate the information entropy of germination *sensu stricto* to the gene expression codifying key enzymes of germination metabolism.

As other molecular techniques, the analysis of gene expression over germination time is used in non-germinated seeds at each point of the water content curve (Rajjou *et al.*, 2011), disregarding impacts of early and/or delayed germinating seeds. However, through information theory, these outliers are capable of altering physicochemical aspects of a developing environment. This can be related to different aspects, such as modifications in water uptake, presence of volatile (mainly VOCs and/or volatile hormones, such as ethylene and jasmonates) and substrate structure (Matilla, 2000; Linkies and Leubner-Metzger, 2012; Motsa *et al.*, 2017; Maurya *et al.*, 2019). In this view, it is expected that the higher the number of random events in the seed-seedling transition system, the greater the information entropy generated (Ribeiro-Oliveira and Ranal, 2016). Thus, findings in seed-seedling transition systems not only can shed light on the understanding of the functional association between information entropy and the early plant development, but also help produce reliable models of theoretical biological systems, an incipient topic in seed science. Therefore, our aim is to promote functional tools for studies on intra-specific variability during the seed-seedling transition.

## Material and Methods

### Biological Model

We used seeds of *Solanum lycocarpum* A. St. ◻ Hil., a nurse plant species from Neotropical areas, as a biological model. This species has been used to understand aspects of the seed-environment interface. The germination transcriptome of this native species presents gene expression of actin, cyclin, heat shock protein, glutathione-S-transferase, malate dehydrogenase and alcohol dehydrogenase similar to congener species such as tomato (Silveira *et al.*, 2019; Souza *et al.*, 2020). What draws our attention in this biological model, however, is the fact that the species presents high intra-specific variability among the seeds that compose the sample, which is the basis of the discussion of the information theory (Shannon, 1948; Labouriau, 1983). The seeds were extracted from ripe fruits collected in 20 mother plants established at 21 ° 12 ’ S, 45 ° 00 ’ W, at 919 meters. The area is a Cerrado (biodiversity hotspot) environment. These seeds were homogenized by the black colour of their tegument and size (4.3 ± 0.3 mm × 5.0 ± 1.0 mm x 0.3 ± 0.1 mm, mean ± SD), with those that were visually damaged, empty or immature being discarded.

### Germination sensu stricto and immediate post-germination

The sowing was performed in plastic germination boxes (gerboxes) on blotting paper. The seeds were equidistantly placed, with the hillum forming a 90° angle in relation to the paper. The volume of water used to moisten the paper was 2.5 times its dry mass. The incubation temperature was alternated between 20 – 30 °C and the photoperiod was of 12 hours (12 h light/12 h Dark) (Pinto *et al.*, 2007).

We calculated weighted mass of germinating seeds (Ribeiro-Oliveira & Ranal 2017 sense), cumulative germination (protrusion of embryo) curves and relative frequency of germination. In addition, the proportion of seed individuals and seedling individuals was also determined over experimental time. Seed germination criterion was embryo protrusion (any embryo structure from teguments). The criterion for determining seedling individuals was developed normal structures for eudicot species, according to ISTA (2017). After reaching this seedling stage, individuals were removed from the germination box in order to not have any potential influence on other germinating seeds, either by volatile emission or by any other component related to embryonic growth, such as anaerobiosis or reduction of water potential due to exudates.

To calculate weighted mass, individual mass of non-germinated seeds was recorded, from the hygroscopic equilibrium (initial seed mass or seed dry mass) to the embryo protrusion. In this case, we analysed 50 seeds (same sample size for each time recorded). In order to promote early plant development phase changes, we applied a differential calculus technique on the weighted mass curve. This calculation demonstrates abrupt changes in the first derivative of the weighted mass curve, i.e. changes in the acceleration due to the water dynamics in germinating seeds (Ribeiro- Oliveira, 2015). It is important to note that we used weighted mass curve as physiological marker, since it was similar as to germinating seeds and young plant in immediate post-germination. Therefore, we prefer to use the most conventional marker to promote the early plant development phase changes. On the other hand, seed germination was characterized using: germinability (G, % embryo protrusion), mean germination time (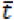; days), mean germination rate (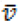; days^−1^) (Labouriau, 1983), coefficient of variation of the germination time (***CV***_***t***_; %), uncertainty (*U*; Bits) and synchronization index (*Z*) (Ranal and Santana, 2006). In this case, we used 200 seeds as a sample (*n*), which were divided in four replicates (*r*), containing 50 seeds each [sub-samples (*ss*); *n* = 200; *r* = 4; *ss* = 50]. Embryo protrusion assessments were made every 24 hours, from seed contact with water up to 15 days after sowing. The end of the assessments took place 16 days after sowing, due to literature protocols (Silveira *et al.*, 2019) on the species and null protrusion in the last observations. The non-germinated seeds had a viable embryo, detected by staining by solution of 2,3,5-triphenyl tetrazolium chloride solution (TTC at 0.1%) with incubation at 30 °C for 24 hours.

### Information entropy for the seed-seedling transition system

The mass data collected over the germination *sensu stricto* time were used to calculate the weighted mass according to Ribeiro-Oliveira & Ranal (2017). From this, the information entropy associated with the seed-seedling transition system was calculated using the following algebraical expression:

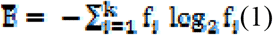

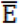: Information entropy associated with seeds germinating in a seed-seedling transition system (Bits); **f**_**i**_: Frequency of water dynamics in germinating seeds, which is defined as:

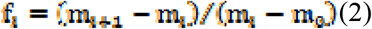

Where: m_i + 1_: water mass in a germinating seed in *i+1* time; m_i_: water mass in a germinating seed in *i* time; m_0_: seed initial mass (dry mass or mass in hygroscopic equilibrium at 25 ± 2.5 °C and 35 ± 2.75% RH).

### Molecular biology studies

We tested our hypothesis by means of molecular aspects related to cell growth and protection as well as the energy metabolism of the embryo in germinating seeds. For that, we used nucleotide sequences proved be efficient for our model species (Silveira *et al.*, 2019).

### RNA extraction and cDNA synthesis

We followed the kit protocols for RNA extraction. Thus, three biological replicates of 50 embryos (*n* = 150, *r* = 3, *ss* = 50) were dissected from germinating seeds by cutting the endosperm with a razor blade. It is important to note that the seeds had been imbibed in water. The dissected embryos were frozen in liquid nitrogen and ground to a powder with the aid of a mortar and pestle in the presence of liquid nitrogen and stored at −80 °C.

We used the commercial NucleoSpin RNA Plant kit^®^ (Macherey–Nagel, Bethlehem PA, USA) for total RNA extraction. Consequently, approximately 100 mg of the ground powder was added to 350 μL of the extraction buffer (RA1) and 3.5 μL de β-mercaptoethanol (β-ME) and then homogenized by vortexing. These samples were transferred to the NucleoSpin Filter^®^ and centrifuged (11 000 rpm) to reduce viscosity. We quantified the extracted RNA using a Nanodrop-2000 spectrophotometer (Thermo Scientific, Wilmington DE, USA) and RNA integrity was confirmed in a 1 % agarose gel (0.4 g agarose + 40 μL de TAE 1X) stained with red gel. The protocol for gel electrophoresis was defined in pretesting and consisted of the running of a solution (500 ng of RNA extracted+ a drop of bromophenol blue) over one hour at 90 Volts.

The commercial kit High Capacity cDNA Reverse Transcription (Applied Biosystems, Foster City CA, USA) was used to perform the cDNA synthesis from RNA of embryos in germinating seeds 1, 5, 10 and 15 days after sowing. As mentioned above, these recorded times were defined by means of a differential calculus applied on the weighted mass curve.

For each biological sample, we used 1 000 ng of total RNA (RT) + 0,8 μL of 25X dNTP Mix (100 mM) + 2 μL 10X RT of random primers + 2 μL of 10x RT Buffer + 1 μL MultiScribeTM Reverse Transcriptase and 4,2 μL Nuclease-free H_2_O. The samples were incubated at 70 °C for 5 min and dipped in ice. The samples were placed in a thermocycler at a temperature of 25 °C for 10 min, 37 °C for 2 h and 85 °C for 5 min for inactivation of the enzymes. The cDNA was quantified in a Nanodrop-2000 spectrophotometer (Thermo Scientific, Wilmington DE, USA).

### Primer design

We designed primers from cDNA sequences of ACTIN (<u>XM-004249818.1</u>), CYCLIN-DEPENDENT KINASE (<u>NM-001247447.1</u>), HEAT SHOCK PROTEIN (SGN-U578134), GLUTATHIONE S-TRANSFERASE (<u>NM-001247228.2</u>), MALATE DEHYDROGENASE (<u>XM-004235934.1</u>) and ALCOHOL DEHYDROGENASE (<u>NM-001247170.1</u>) that are available from the NCBI database (http://www.ncbi.nlm.nih.gov/). This was done according to Silveira and Ribeiro-Oliveira et al. (2019). We also used the amino acid sequence of METALLOTHIONEIN (SGN-U143485) to design the primers. This gene was chosen as a reference gene since it presents Ct similarity over seed germination time (Ct between 18.90 and 18.95). The amino acid sequence was converted into a nucleotide sequence using the tool PerlPrimer (http://perlprimer.sourceforge.net; Marshall 2004). Sequences were then submitted to Blast (https://blast.ncbi.nlm.nih.gov/Blast.cgi) to confirm the primers (Table 1–2).

**Table 1.**
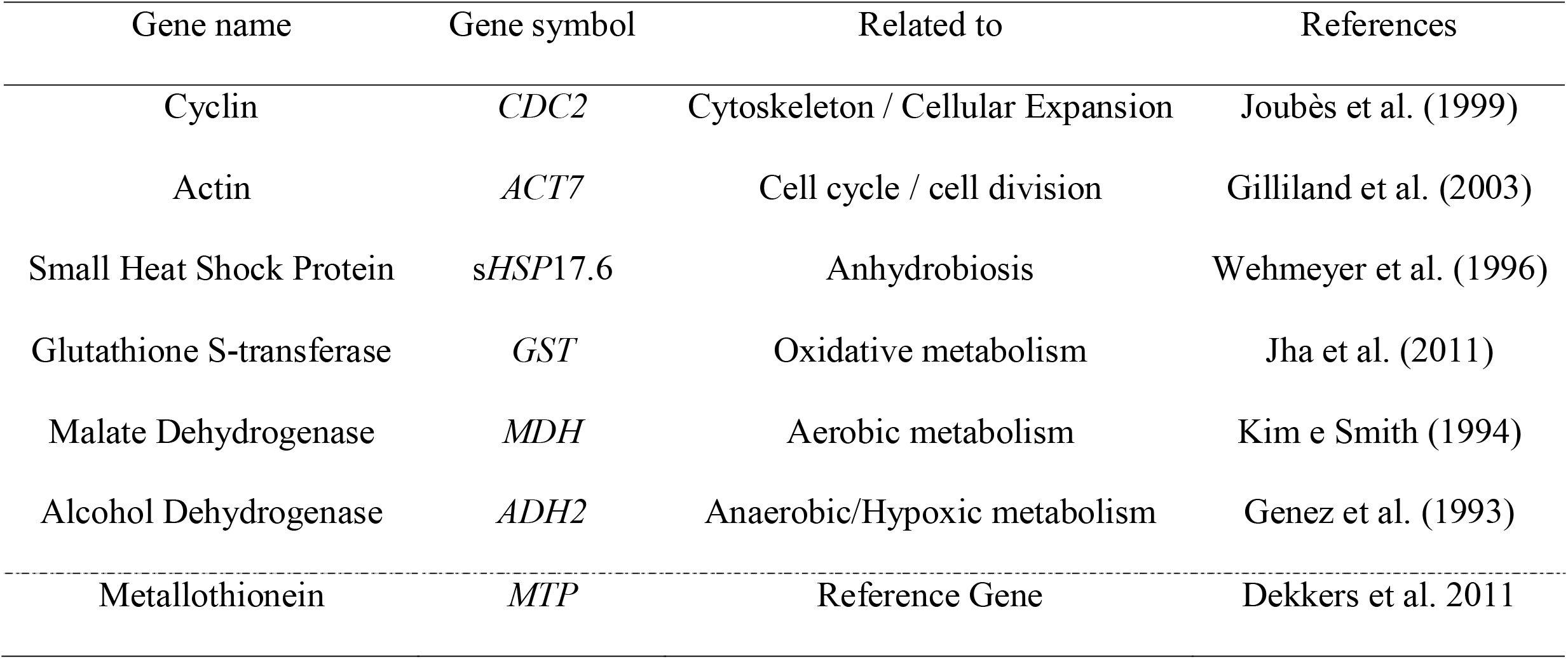
Genes (mRNAs) studied for the embryo development in germinating seeds of *Solanum lycocarpum* A. St.-Hil.

| Gene name | Gene symbol | Related to | References |
| --- | --- | --- | --- |
| Cyclin | *CDC2* | Cytoskeleton / Cellular Expansion | Joubès et al. (1999) |
| Actin | *ACT7* | Cell cycle / cell division | Gilliland et al. (2003) |
| Small Heat Shock Protein | *sHSP17.6* | Anhydrobiosis | Wehmeyer et al. (1996) |
| Glutathione S-transferase | *GST* | Oxidative metabolism | Jha et al. (2011) |
| Malate Dehydrogenase | *MDH* | Aerobic metabolism | Kim e Smith (1994) |
| Alcohol Dehydrogenase | *ADH2* | Anaerobic/Hypoxic metabolism | Genez et al. (1993) |
| Metallothionein | *MTP* | Reference Gene | Dekkers et al. 2011 |

**Table 2.**
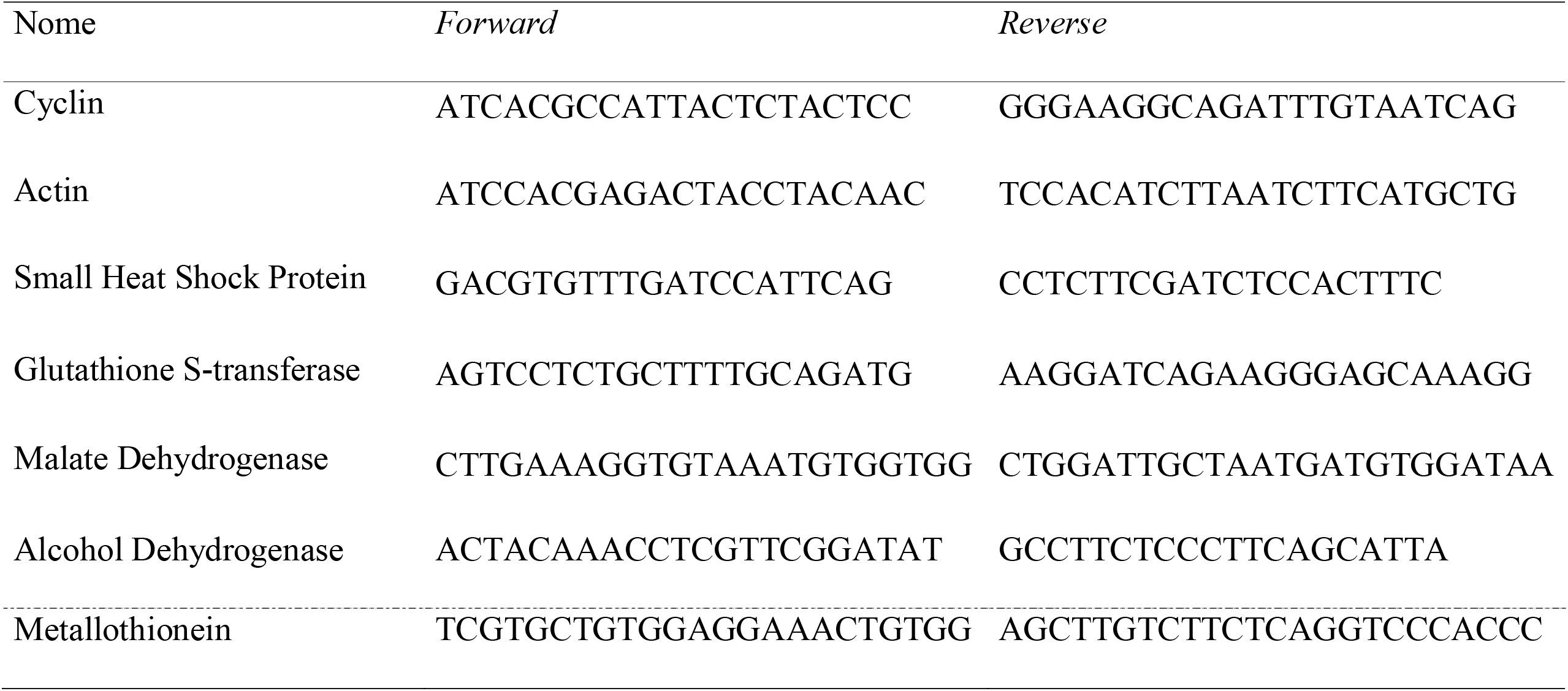
Primers for real-time PCR analysis of embryo development in germinating seeds of *Solanum lycocarpum* A. St.-Hil.

### Statistical analysis

First, we submitted all the data set (germination *sensu stricto*, post-germination and molecular raw data) to the bootstrap method with 1 000 re-samplings, since values generated above this number are similar according to the convergence test. This was an assumption to calculate weighted mass curves and then, means and curves of information entropy. We used bootstrap because it is a technique of successive resampling from original data (Efron, 1979). This ensures that analytical models are reproducible and reliable. In addition, the bootstrap technique is useful in applications for which analytical confidence intervals are unobtainable or when robust nonparametric confidence intervals are required (Buckland, 1985). It also estimates the distribution of an estimator, reduces impacts of outlier and numerical anomalies, calculates the estimates of standard error and the population parameters of confidence intervals (IBM, 2015). To calculate these confidence intervals (α = 0.05), we used the Algorithm AS 214 for Fortran (Press *et al.*, 1992), which is recommended to perform the Monte Carlo Confidence Intervals.

For molecular comparisons of fold change, we used the tool REST® for Ct calculation, which was based on the data simulation of bootstrap methods from raw data. The REST performs comparative quantifying from the Pair-Wise Fixed Reallocation Randomization Test method (Pfaffl, 2002) using a normalizing gene.

Finally, we performed the correlation test for residuals of characters calculated both in the physiological and in the molecular analyses by means of pair-wise Spearman rank correlation (*P* ≤ 0.01). From the pair-wise Spearman correlation coefficients we generated the correlation matrix heatmap according to Heatmapper software (Babicki *et al.*, 2016).

## Results

The differential calculation technique applied on the weighted mass curve detected four phases of the seed-seedling transition process (germination *sensu stricto* and early development of young plants in post-germination) of *Solanum lycocarpum* (Fig. 1). The most stable phase began approximately 12 hours after seed contact with water, when seed imbibition ended. The first germination occurred approximately 9 DAS, when the third seed-seedling transition phase started; this is the moment that the number of young plants in immediate post-germination increases. From that moment on, however, the increase in the proportion of seedling individuals in relation to the seed individuals was gradual. This proportion was effectively greater after the fourth seed-seedling transition phase started. It is important to note that seeds had around 74% germinability and slow kinetics 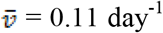 for embryo protrusion. This protrusion was a quite uniform process over time (CV_t_ = 20.01%). Additionally, this process was not very predictable (U = 2.27 bits) nor synchronous (*Z* = 0.25) showing high intra-specific variability (Fig. 1).

**Fig. 1:**
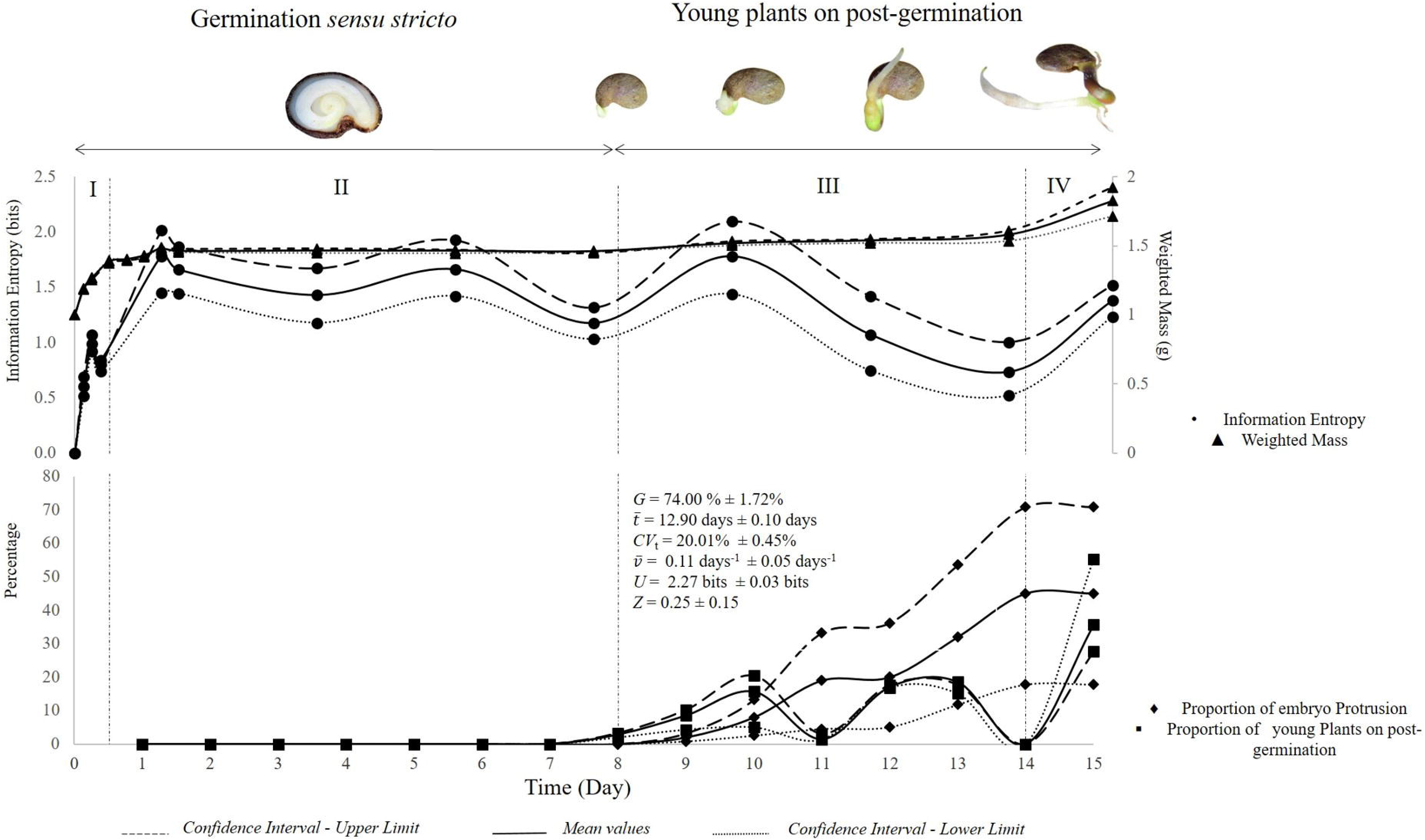
Seed-seedling transition in *Solanum lycocarpum* A. St.◻Hil. Development phases (see vertical black line) were defined from a calculation technique applied on the weighted mass curve (upper secondary axis), which was determined by the velocity of water dynamics on germinating seed. Information entropy (upper primary axis) is related to water dynamics in germinating seeds in seed-seedling transition system. Percentage axis describes relative frequencies of embryo protrusions and seedling development. In all cases, dashed lines represent lower and upper confidence intervals; while filled lines represents mean values (1 000 Monte Carlo simulations at 0.05). Values of classical germination measurements (mean ± confidence intervals) were also obtained from 1 000 Monte Carlo simulations at 0.05. *G*: Germinability (%); ***t̄***: mean germination time; *CV*_t_: coefficient of variation of the germination time; ***v̄***: mean germination rate; *U*: Uncertainty; *Z*: Synchronization index.

The end of anhydrobiosis, marked by the metabolic intensification resulting from the entry of water into seed tissues, still in phase I, caused a fast increment of information entropy (Fig. 1). One of the highest information entropy values of germination *sensu stricto* in the middle of this phase indicated the end of imbibition per se and the beginning of a predominantly biophysical stage, in which although biophysical process are predominant, biochemical processes also act. The alteration of the predominantly biophysical phase to the first predominantly biochemical phase (phase II) is marked by a small reduction in information entropy. This phase is characterized by the up-regulated of genes codifying Alcohol dehydrogenase in relation to the down-regulated of genes codifying Malate dehydrogenase (Fig. 2). In addition, in phase III, there is also up-regulated of genes codifying protective enzymes, GST and HSP, as well as growth related proteins, as actin. Although down-regulated, genes codifying cyclin are still expressed in a significant way during phase II (Fig. 2).

**Fig. 2:**
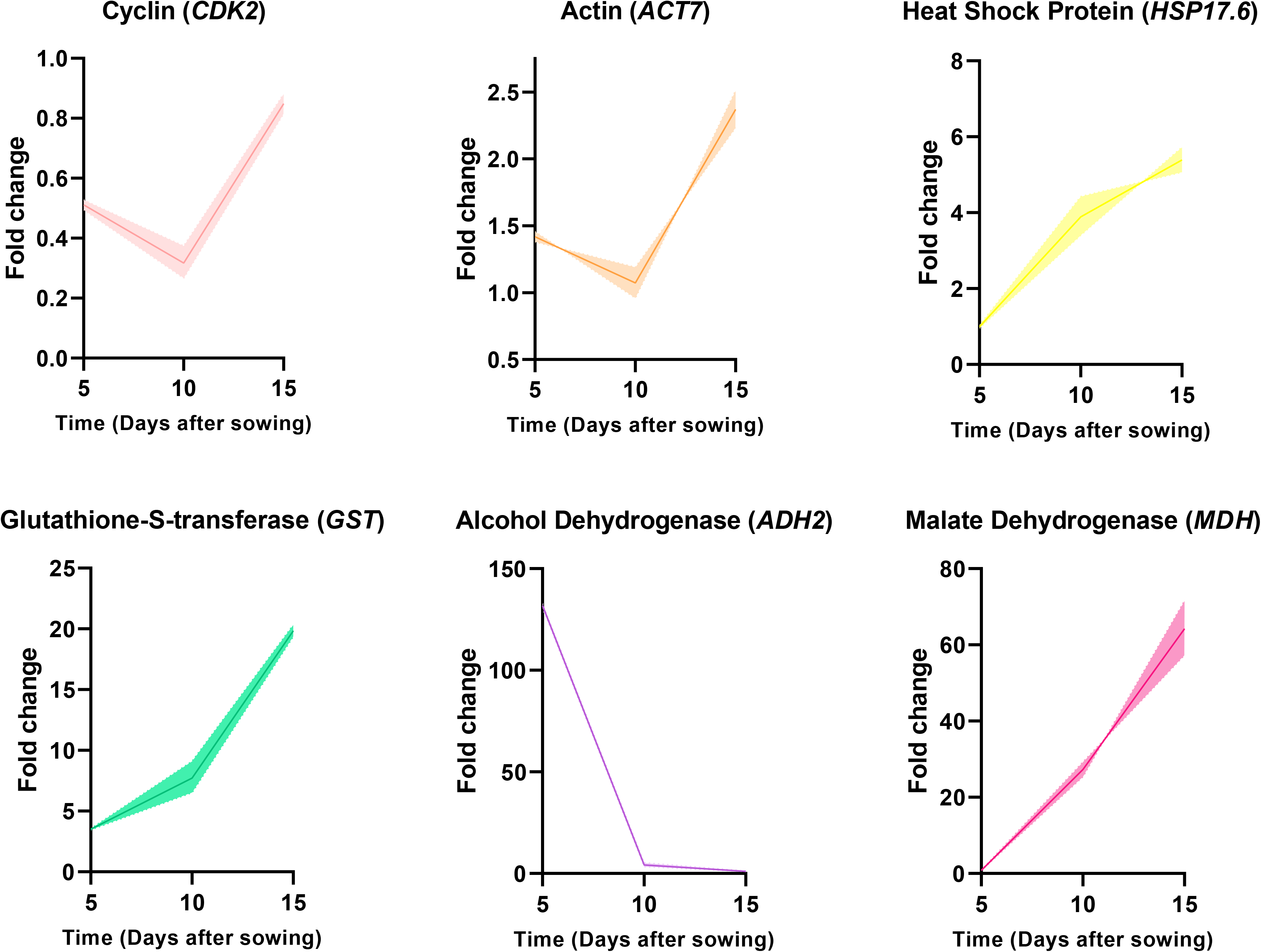
Relative fold change in transcript abundance of genes related to cellular division (Cyclin – CDK2) and expansion (Actin – ACT7), anhydrobiosis tolerance (sHSPs – HSP17.6), oxidative metabolism (GST – GST) and respiratory metabolism (anaerobic: MDH – MDH – and anaerobic: ADH – ADH2) of embryos in germinating seeds of *Solanum lycocarpum* A. St.-Hil. imbibed in water. The error bars indicate interval confidence (α = 0.05) and mean values (filled line) are based on the bootstrap method (1 000 resampling) applied to raw data. For each biological triplicate were used 50 embryos (*n* = 150; *r* = 3; *ss* = 50).

The first protrusion reduces information entropy quickly, which coincides with the transition to the third stage of the seed-seedling transition (Fig. 1). At this stage, the only peak of information entropy is 10 DAS, a moment prior to the peak of embryo protrusion in the sample 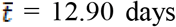. In this peak, there are increments of gene expression (up-regulated) related to aerobic and protective respiratory metabolisms; while there is a reduction in gene expression codifying enzymes involved in embryo development *per se* (actin; up-regulated, and cyclin; down-regulated) and the anaerobic metabolism (ADH) (Fig. 2). The increase in the proportion of seedling individuals in relation to the seed individuals is accompanied by a decrease of information entropy, which reaches the lowest value when the proportion of seedling individuals becomes larger than the seed Individuals (Fig. 1). This is also the moment the last stage of the seed-seedling transition starts, characterized by the higher proportion of seedling individuals in relation to the seed individuals in *Solanum lycocarpum*. This demonstrates the efficiency of the measurement, which is not affected by the increase of water mass at the beginning of the establishment of young plants. The highest proportion of individuals in the same physiological phase increases the information entropy of the seed-seedling transition system. This is accompanied by the reduction of genes codifying ADH (here down-regulated) and the increment of genes codifying protective molecules (GST and HSP), aerobic respiration (MDH) and cell development (actin and cyclin), which are up-regulated.

In general, information entropy is negatively correlated to the seed-seedling transition (Fig. 3). When we evaluated each development phase, however, we noted the positive linear correlation between development measurements and information entropy, demonstrating that development peaks increase the entropy of the seed-seedling transition system. On the other hand, information entropy is negatively correlated (linearly) with mRNA codifying cellular protective enzymes (Fig. 3). What is noteworthy is the fact that the progress of the development phases denotes subtle differences in individuals, from the physiological point of view. These differences are most evident in the seeds that germinated very early or late in the sample (outliers). Since the number of outliers is small, the communication between seed-individuals is also smaller, which causes the communication disturbance to be lower and, consequently, the information entropy is reduced.

**Fig. 3:**
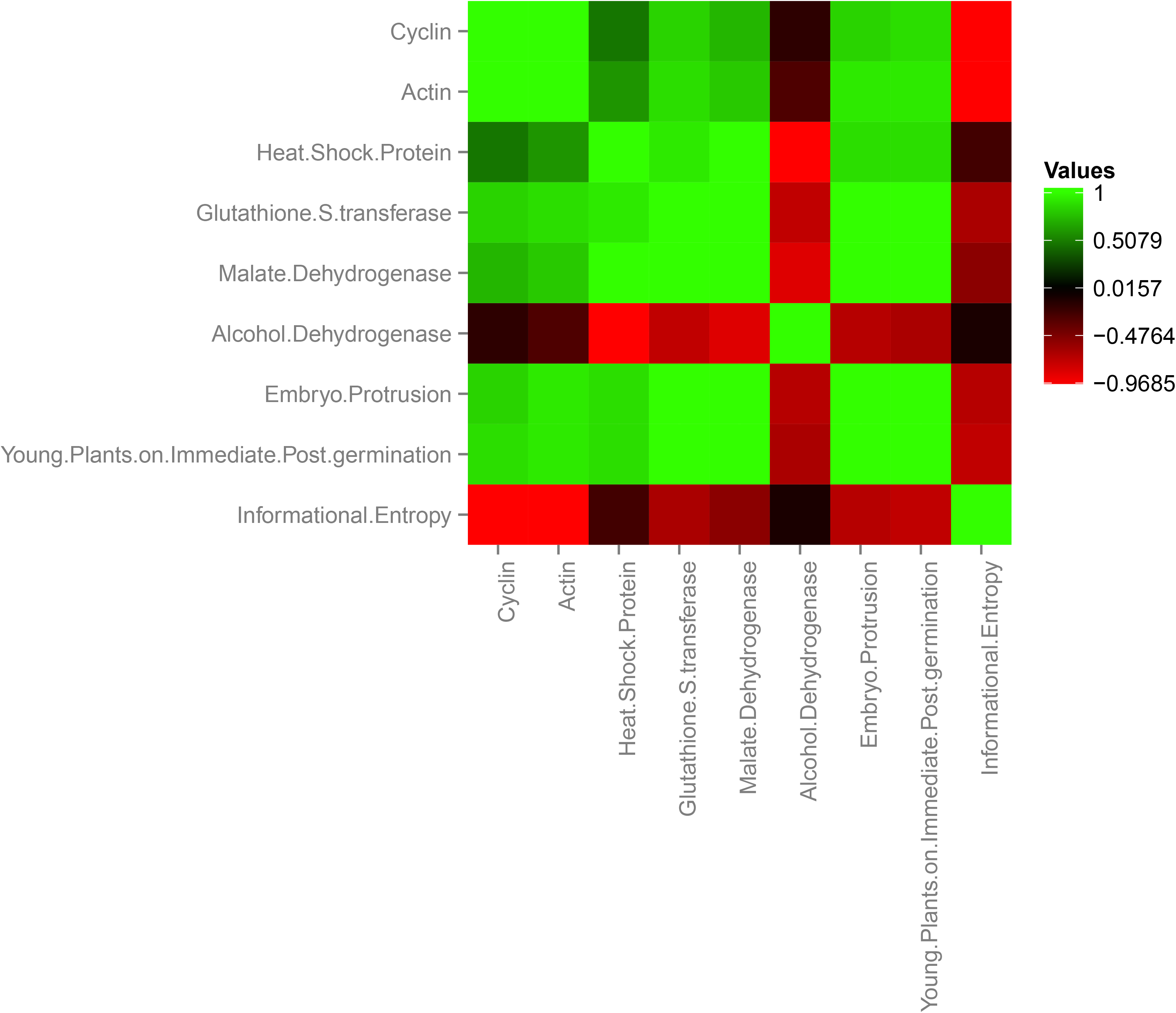
Correlation matrix heatmap based on pair-wise Spearman correlation coefficients (rank correlation; *P* ≤ 0.01) for physiological and in the molecular analyses.

## Discussion

We demonstrate that information entropy in a seed-seedling transition system has an influence on the number of individuals in the sample in each physiological stage (seed or seedling). At first, this could be considered a consequence of the measurement being influenced by sample fluctuation (Peet 1975; Wolda 1981; Ribeiro-Oliveira, Ranal & Santana 2013), but this was not the case, since the calculations for the measurement were performed for the same sample size. Therefore, it is possible that germinating seeds and/or seedlings present a sophisticated intra-specific communicative channel that guides embryo protrusion and, consequently, post-germination.

Here, entropy is the disturbance generated in the seed-seedling transition system by adhesion of water molecules to the seed. Thus, the greater the entropy generated by water, the lower the uniformity of absorption, related to the amphibolic process that nourishes the embryonic growth/development [see details on germination amphybolism in Bewley, Bradford, Hilhorst & Nonogaki (2013)]. This becomes more evident when one takes into account that the metabolic intensification of germination arises with the adhesion of the water molecules to the seeds, generating a complex and extremely dynamic metabolic trigger, which causes great movement (energy) in the beginning of the predominantly biochemical phase (phase II). It is noteworthy that entropy (a term used for the first time by the German physicist Rudolf Julius Emmanuel Clausius, in 1865) is a physical magnitude that, in thermodynamics, measures the degree of irreversibility of a system (Schroeder and Pribram, 1999; Leff, 2007; Frigg and Werndl, 2011), which is consistent with embryo development. Entropy demonstrates how matter and energy are stored and distributed across the boundaries of the thermodynamic system (Frigg and Werndl, 2011). Thus, although there is a clear distinction between the magnitudes of mass, internal energy and quantity of matter, the entropy of a system is certainly related to these, therefore it is also a property of the system (Leff, 2007; Frigg and Werndl, 2011). There is a tendency of nature to equalize processes as order passes into disorder, that is, when molecular disorder increases. Thus, Ludwig Boltzmannn, applying the concept to physics, in 1896, stated that entropy can be defined as a measure of the degree of system disorder (Leff, 2007; Haynie, 2008; Dragicevic and Sredojevic, 2011). This explains the close relationship between entropy and embryo development under a standard temperature, as was the case with the experimental environment. It is important to emphasize that what is called entropy here is the contribution of mass to thermodynamic entropy, which is conceptually information entropy.

By transporting the thermodynamic knowledge to seed science and with the revisiting of Shannon (1948) on entropy, Labouriau & Valadares (1976) defined the information entropy index of the distribution of relative frequencies of germination on the variation of the liquid enthalpy of activation. These indexes, calculated in order to discover the optimum temperatures for the germination of seeds, follow the assumption that the seed is a thermodynamic system. Intuitively, these authors related two aspects that, according to modern physics, may be intimately linked. Shannon's information theory, in turn, says that entropy keeps close relationship with the state of information of a system. Entropy acts in the sense of always measuring the loss of information of the system. Given this, the greater the information, the greater the entropy (Shannon, 1948; Jost, 2006). This justifies the maximum entropy in the seed-seedling transition occurring when all the cells are in operation and with most of the individuals in the system at the same physiological stage. This occurs after the predominantly biochemical phase starts, at the peak of embryo protrusion and at the peak of post-germination development.

Entropy is as important for biological systems as energy and it influences life in a similar way; however, unlike energy, that can be retained, entropy cannot and, in a natural process, always increases and this is how we know if the process is spontaneous (Schroeder and Pribram, 1999; Leff, 2007; Frigg and Werndl, 2011). Taking this into consideration, our findings show that the germinative process is only spontaneous in the imbibition phase per se. From this, there is the performance of biochemical complexes that manage the cellular turgor and, consequently, the biases of this process. Thus, the maximum entropy points of the seed-seedling transition system characterize moments in which there is the maximum expression of intra-specific variability, in which the germinative pattern of the seed stands out individually in relation to the sample pattern. This means that in these moments it is possible to denote individual differences (intra-specific variability) regarding the perception and reaction to environmental clues. Therefore, moments of maximum entropy in the seed-seedling transition system can be crucial points for the decision of the seed to germinate or maintain anhydrobiosis.

The germination of *Solanum lycocarpum* seeds occurs in a cadenced way (see low ***v̄***), over days, due to the slow activation of enzymes related to the metabolism of structural carbohydrates (the weakening of endosperm by endo-beta-mannanase according to Pinto *et al.,* 2007). This leads to a gradual and inversely proportional change from an anaerobic to an aerobic metabolism in the embryo, something expected and which supports the gradual embryo development. This ensures that the linkage between information entropy and change of transcripts becomes more assertive, decreasing the probability that the analysis time may disregard important aspects of the metabolism. In this way, it is important to emphasize that 74% of the seeds of the studied sample germinated, but those that did not germinate at the end of data collection were viable. This demonstrates that all seeds in the sample had biological activity in the system. Moreover, the closure of the analysis process was made in resonance with other studies of molecular and physiological aspects of the germination of the species, whose seeds are anhydrobiotic (= orthodox) and non-dormant (Silveira *et al.*, 2019; Souza *et al.*, 2020).

The unquestionable correlation between the information entropy peaks and the gene expression codifying important enzymes for the germination metabolism reinforces the assertiveness of our hypothesis. In this sense, one approach is that the relationship between information entropy and growth processes is a very concrete aspect that can assist in the production of synthetic biological models for seed science. What draws more attention is the fact that the seed individual and seedling individual proportions are determinant for the genetic activities of the embryo. When the proportion of individuals in the sample was predominantly of seeds, the information entropy was greater, as well as the expression of genes codifying key enzymes for embryonic development; When the proportion of seedlings was higher, even with all the seeds of the sample still viable, the expression of these genes in the embryo was smaller, conferring less information entropy in the system. These two facts demonstrate that (i) according to physics, mathematics and ecology reports, information entropy enables the quantification of disturbance in the system relative to individuals in the same physiological stage (seed-seed or seedling-seedling), which may be determinant for intra-seminal embryonic development. (ii) There is a possible intra-specific communication in systems formed by germinating seeds with the potential to alter the pattern of embryonic development of the sample. These aspects, which have long been discussed by ecophysiologists (e.g. Peet, 1975), may come from physiological processes associated with maternal recruitment, as well as from the seed, such as volatile emission (Ribeiro-Oliveira and Ranal, 2016). In either way, there are still few reports that decipher how this effect of mutual stimulation promotes the germination boom, and information entropy can stand out as a useful measurement for this. Therefore, we suggest the use of information entropy as a tool for biological systems studies.

## Acknowledgements

We are grateful to the Programa de Pós- Graduacão em Agronomia UFU (PPGA-UFU) and Coordenação de Aperfeiçoamento de Pessoal de Nível Superior(CAPES, Financial code 001) for the support currently given to the first author as a Post-doctoral (PNPD); and to Mr. Roger Hutchings for the English review of the manuscript.

## Author contributions

JPR-O planned experiments, analysed and interpreted the data, and wrote the manuscript; LEDS and LAVP performed experiments; HWMH and EAAS contributed equally, supervising and contributing to the experimental data collection and manuscript preparation.

